# Responses of Agricultural plants to Lithium pollution: Trends, Meta-Analysis, and Perspectives

**DOI:** 10.1101/2022.05.07.491047

**Authors:** Noman Shakoor, Muhammad Adeel, Imran Azeem, Muhammad Arslan Ahmad, Muhammad Zain, Aown Abbas, Pingfan Zhou, Yuanbo Li, Xu Ming, Yukui Rui

## Abstract

Lithium (Li) is gaining attention due to rapid rise of modern industries but their ultimate fingerprints on plants are not well established. Herein, we executed a meta-analysis of the existing recent literature investigating the impact of Li sources and levels on plant species under different growth conditions to understand the existing state of knowledge. Toxic effects of Li exposure in plants varies as a function of medium and interestingly, more negative responses are reported in hydroponic media as compared to soil and foliar application. Additionally, toxic effects of Li vary with Li source materials and LiCl more negatively affected plant development parameters such as plant germination (n=48) and root biomass (n=57) and recorded highly uptake in plants (n=78), while LiNO_3_ has more negative effects on shoot biomass. The Li at <50 mg L^-1^ concentrations significantly influenced the plant physiological indicators including plant germination and root biomass, while 50-500 mg L^-1^ Li concentration influence the biochemical parameters. The uptake potential of Li is dose dependent and their translocation/bioaccumulation remains unknown. Future work should include complete lifespan studies of the crop to elucidate the bioaccumulation of Li in edible tissues and to investigate possible trophic transfer of Li.

**Environmental significance:** Accumulation of Li sources is increasing in ecosystem compartments, and this might be vulnerable to plants.

## 1. Introduction

Lithium (Li) is the major element powering modern technology known as white gold and a green energy alkali metal ^1^. Globally, total Li production was 82,000 tons year^−1^ in 2020, a 200% increase since 2010 (USGS 2011-2021). Rechargeable Li-ion batteries consumption was 74% of total Li production by 2021^2^, which shows the increasing demand of Li for electric vehicles and products. ^3^. Soaring demand for Li had upregulated the global supply of Li exponentially to maintain the technology progress, at the same time electronic waste is significantly contributing to the contamination of soil with Li^+^ ions or Li_2_O.^4^ Lubricating grease contains 4% of total Li industrial application ^5^, which might enters the environment through runoff from roads. The production of glass and ceramics consumes 14% of Li total production,^6^ which finally goes to landfill. In addition, antidepressants drugs contain Li which ultimately enters to the soil system through landfill leachate, sewage and rainwater runoff.^7, 8^

Globally, extremely volatile concentrations of Li in soil, water and in agricultural ecosystems have led to fluctuations in Li concentration in plants, food products posing risk to humans.^9, 10^ Recently, different studies reported the effects of Li on plants and found a wide variety of harmful, neutral and positive effects on plants.^11-13^ Despite the possible entry of Li in agricultural ecosystem, understandings on interaction of Li in agricultural crops is not yet clear, like in our scientific survey we observed that studies reported contradictory findings about Li exposure to agricultural crops.^14, 15^ Previous studies such as, Jiang et al. ^14^, Li et al. ^16^ and Li, Guo et al ^17^ performed a detailed investigation of Li exposure to *Apocynum venetum L, Brassica carinata*, and *Apocynum pictum*. Jiang et al. treated *Apocynum venetum L* plants with LiCl for 90 days, and found that plant dry weight, chlorophyll content and stomatal conductance was notably reduced with increasing concentrations. However, Li accumulation in leaves was high at 200 mg kg^-1^ as compared to 50 mg kg^-1^ and start decreasing beyond this level.^14^ Similarly, Li et al. found that when *Brassica carinata* was exposed to LiCl at different concentration (0.03-120 mM) in petri dishes, a markable reduction in germination rate, root length of seedling and fresh weight was noticed at high concentration. The increase in growth rate was observed only at 0.03-30 mM concentration level. Interestingly, a significant decrease was observed in chlorophyll content at high concentrations which inhibited the growth of the *Brassica carinata* seedling. ^16^ Bakhat et al found the spinach shoot and root dry biomass significantly increased with low level of Li exposure.^18^

In contrast, Jiang et al. conducted the same experiment with another plant species namely *Apocynum pictum*, and documented that LiCl (25 mg L^-1^) increased the germination percentage (>80%) as compared to control and other applied concentrations. Dry weight of plants showed a non-significant effect at 50 mg kg^-1^, but reduced at 200 and 400 mg kg^-1^. A significant decrease in chlorophyll a, b and carotenoids content was also observed at high level (200 and 400 mg kg^-1^).^17^ Moreover, two studies reported the deleterious effects of foliar application of Li on *Lactuca sativa L*. and *Alvia hispanica* plants which were different from soil and hydroponic exposure.^13, 19^ Consequently, it is often challenging to assess the effect of the Li threshold level on agriculture plants. In above mentioned context, it is necessary to evaluate the toxicology of Li for a comprehensive assessment of hazards on agroecosystems.

Given the expected discharge of Li into the environment, we first time established current state of knowledge through the meta-analysis after an intensive look at published literature. Our meta-analysis investigates the importance of Li exposure by its sources, medium and concentration on the plant growth parameters. Through this comprehensive research analysis, we also highlighted research gaps, proposed methods, recommendations and perspectives for future research, and provide an updated baseline for scientists working in this field. Current study provides the way forward for additional research work to mitigate the Li pollution and to accurately assess the fate of Li in agricultural environments.

## 2. Methodology

### 2.1 Literature search

A total of 790 studies that reported the interaction of Li in the ecosystem were collected from Google Scholar, Web of Science, Science direct, PubMed, and other web sites. Keywords used for the search includes “LiCl”, “Li_2_SO_4_”, “LiOH”, “transfer”, “toxicity”, “soil” and “plants”. We have developed and implemented a strict search strategy to collect the novel, highly relevant and reliable data sets (Figure 1). As mentioned above, the primary search returned 790 articles and this group was limited to 58 research articles by making the following criteria mandatory in our search strategy: (a) the experiment was evaluated under proper research plan (field or laboratory condition) (b) the study has included the application of Li on plants (c) all the results were held by appropriate statistical analysis and (d) control treatment was included in study. Only those studies were included for meta-analysis that met all the above criteria.

**Figure 1.**
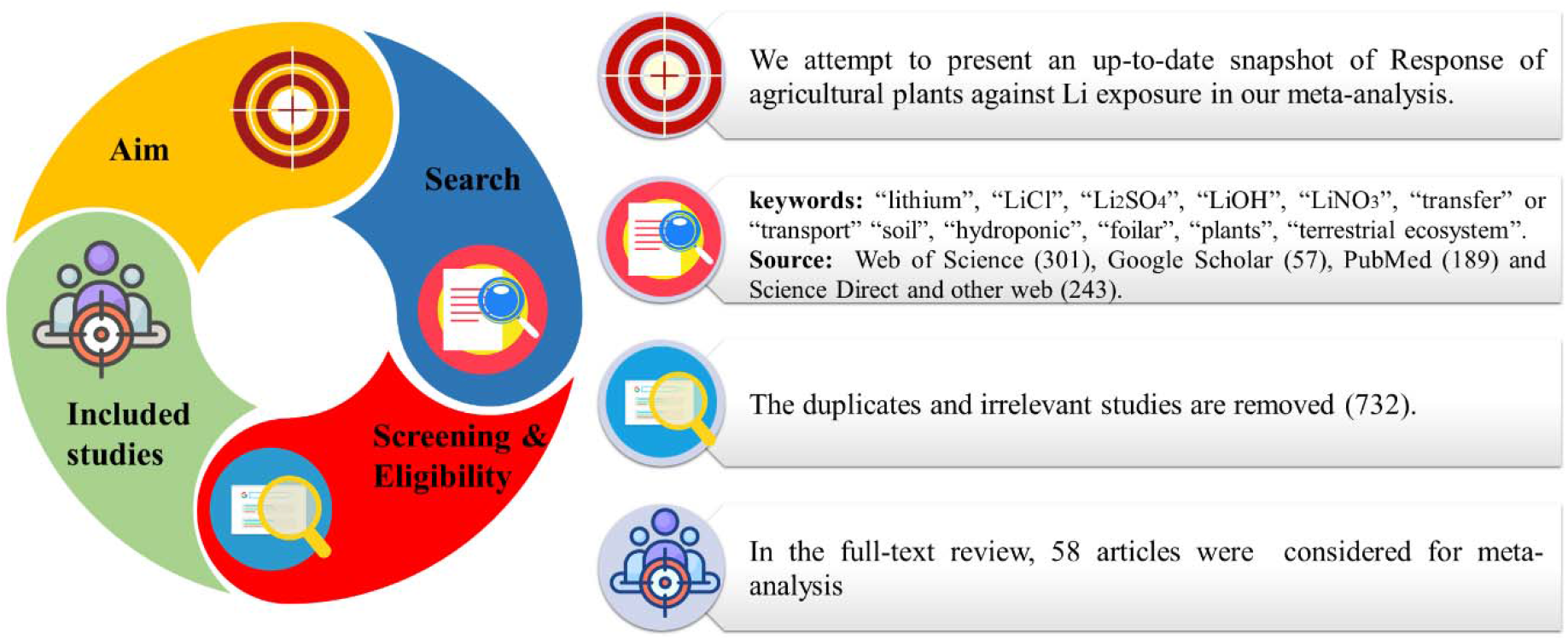
Strategy of work plan for conducting the meta-analysis

### 2.2 Data extraction

Data was obtained from each research study that met all four criteria, together with exposure medium (soil, hydroponic and foliar application), concentration and Li sources [lithium sulfate (Li_2_SO_4_), lithium nitrate (LiNO_3_), lithium chloride (LiCl) and lithium hydroxide (LiOH)]. In the meta-analysis total 58 peer-reviewed research articles were selected; the data was recovered from the table directly. The Get Data Graph Digitizer software (version 2.26) and Web Plot Digitizer software (version 4.5) were used to assemble data from the figures in published research articles.

### 2.3 Data endpoints

In current study categorized all the endpoints (number of each parameter in the studies) into six categories depending upon their biological significance. Biological significance includes germination, shoot length, root length, chlorophyll (a and b) and Li uptake (Figure 2). Furthermore, if the authors of an individual study recognized different observations of Li and studied endpoints such as exposure medium, plant species, Li source, and different concentration, we considered each endpoint separately for each observation. The endpoints of each parameter (overall) are expressed as a percentage.

**Figure 2.**
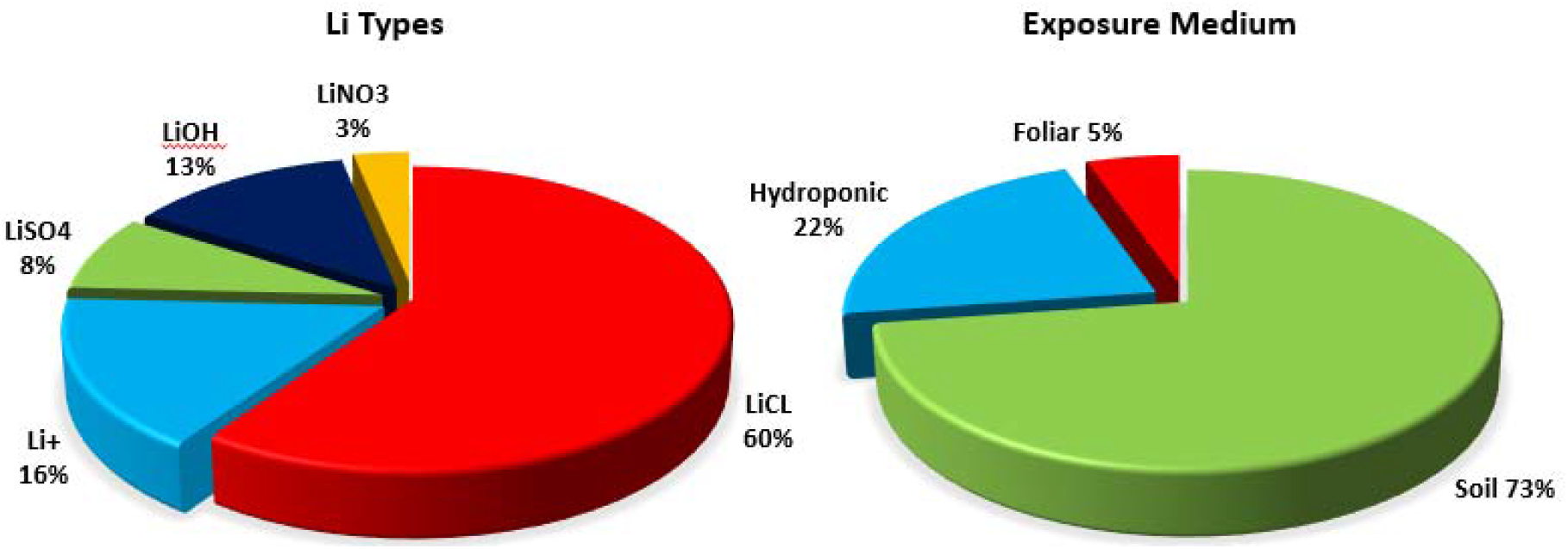
Published literature regarding the effects of Li on plant species. (A) Li source evaluated (B) Medium exposure of Li.

### 2.4 First-order meta-analysis

In current meta-analysis, the natural log-transformed response ratio (lnRR) technique were used as documented in Gurevitch, et al. ^20^ and Hedges, et al. ^21^ to assess the impact of exposure medium, Li source, and concentration against plant germination, shoot length, root length, chlorophyll a, chlorophyll b and Li uptake (Figure 2). The equation used for analyzing the positive, negative or neutral effect is given below.

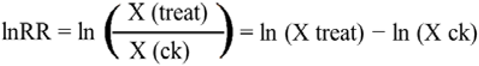

Where X (ck) shows the mean of control treatment, while X (treat) represents the mean of th treatment. The few studies shown the standard error, we calculated standard deviation (SD) from standard error using formula SD = SE √(n) (n is the number of replicates). The estimated size impact was converted into percentage by using the equation given below. (elnRR −1) × 100%

### 2.5 Statistical analysis

All statistical analysis and the data pre-processing were executed in Microsoft Excel 2013, and OpenMEE software which heavily leverages and can largely be considered as a GUI to the metaphor^22^ R packages. The experimental polled effect was based on the first-order meta-analysis with 95% confidence intervals (CI = 95%) and was presented as forest fragments. A positive value shows a positive effect on plant, and negative values shows a negative impact.

## 3. Results and Discussion

### 3.1 Li exposure medium consequences on plants

In most of the endpoints, plants were exposed to Li via soil (48%), while in others exposure through hydroponic and foliar application was 42% and 10% respectively (Figure 3). A negative response on plant germination, root and shoot biomass resulted from Li exposure and this response varied with exposure medium (Figure 3A, B, and C). Under hydroponic exposure, plant germination (n=79), root (n=80) and shoot biomass (n=157) was significantly decreased by 15%, 18% and 20% respectively whereas, soil exposure reduced the corresponding parameters by 8%, 10% and 17% respectively. Furthermore, foliar application of Li increased plant germination (10%) and root biomass (13%), while decreased (8%) the shoot biomass. Average across the exposure medium, overall Li reduced the germination, root and shoot biomass by 5%, 7% and 15% respectively. *Apocynum venetum* exposed to LiCl at 50, 200 and 400 mg kg^-1^ have reduced the root and shoot biomass by 4, 40, and 86% respectively, while reduction in shoot biomass at these values was 20, 52, and 75% respectively under soil medium. ^23^ Bakhat et al.^11^ observed that the exposure of LiCl to spinach increased the root biomass at 20, 40, 60 and 80 mg kg^-1^ by 64, 27, 15, and 12% under soil. However, the reduction in shoot biomass was noticed by 15, 17, and 16% at 40, 60 and 80 mg kg^-1^ respectively except at 20 mg kg^-1^ that increased shoot biomass by 16%.^11^ Application of LiCl to *Zea mays* at 50 mg dm^-1^ under hydroponic medium inhibited the root biomass by 31%. Similarly, shoot biomass of sunflower and *Zea mays* was reduced by 27 and 32% respectively under same concentration and medium.^24^ Interestingly, foliar application of Li significantly enhanced the *Lactuca sativa L*. root biomass by 29 and 41% at 20 mg dm^-1^ and 30 mg dm^-1^ while declined the shoot biomass by 15 and 19% respectively.^13^

**Figure 3.**
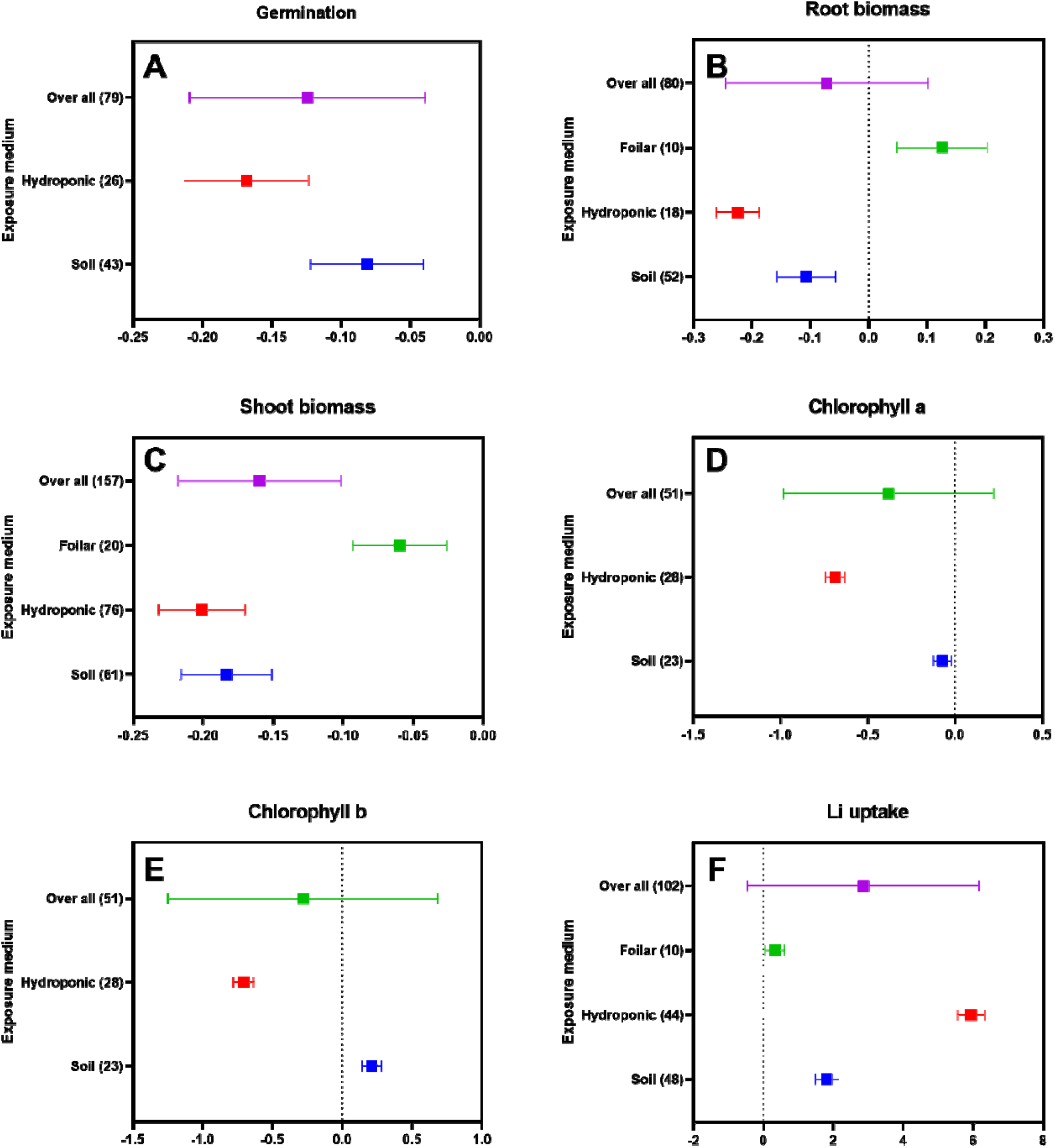
Mean effect of exposure medium on A) germination, B) root biomass, C) shoot biomass, D) chlorophyll a, E) chlorophyll b, and F) Li uptake. The lift side bars highlighted the exposure medium are presented, and the corresponding number of endpoints is presented within the parentheses.

Overall, chlorophyll a and b contents were reduced by 50% and 54% upon Li exposure in hydroponic medium. In contrast, chlorophyll a contents were decreased by 7% and chlorophyll b contents were increased by 23% at soil medium (Figure 3D and E). The previous study revealed that the chlorophyll b increased in the spinach leaves at 40-80 mg kg^-1^ of LiCl.H_2_O as compared to control plant ^18^. Whereas, the exposure of LiCl significantly reduced the chlorophyll a and b contents of *Apocynum venetum* by (22 and 13%), (53 and 50%) and (83 and 88%) at 50, 200 and 400 mg kg^-1^ respectively under soil medium.^14^ Another study reported that chlorophyll contents (a and b) were significantly reduced in the *Brassica carinata* seedling in petri dishes at >60 mM LiCl treatment. ^25^ The chlorophyll a and b content in the *Zea mays* plant significantly reduced by 47-43% with increasing the of Li in hydroponic medium (25 and 50 mg dm^-3^).^12^

Li uptake by plant was also affected under different exposure, as significant increases were observed in order of hydroponic medium > soil medium > foliar application (Figure 4F). Bakhat et al. 2020 revealed that Li exposure to soil significantly enhanced the Li accumulation in shoots and root of spinach in three harvesting.^18^ As the concertation increased the uptake of Li was increased. In short, Li may have negative impact plant germination, root biomass, shoot biomass, chlorophyll a and b in hydroponic condition as with soil medium and foliar application. it may be due to Li ions quickly absorbed by plants and effect the plant growth.

**Figure 4.**
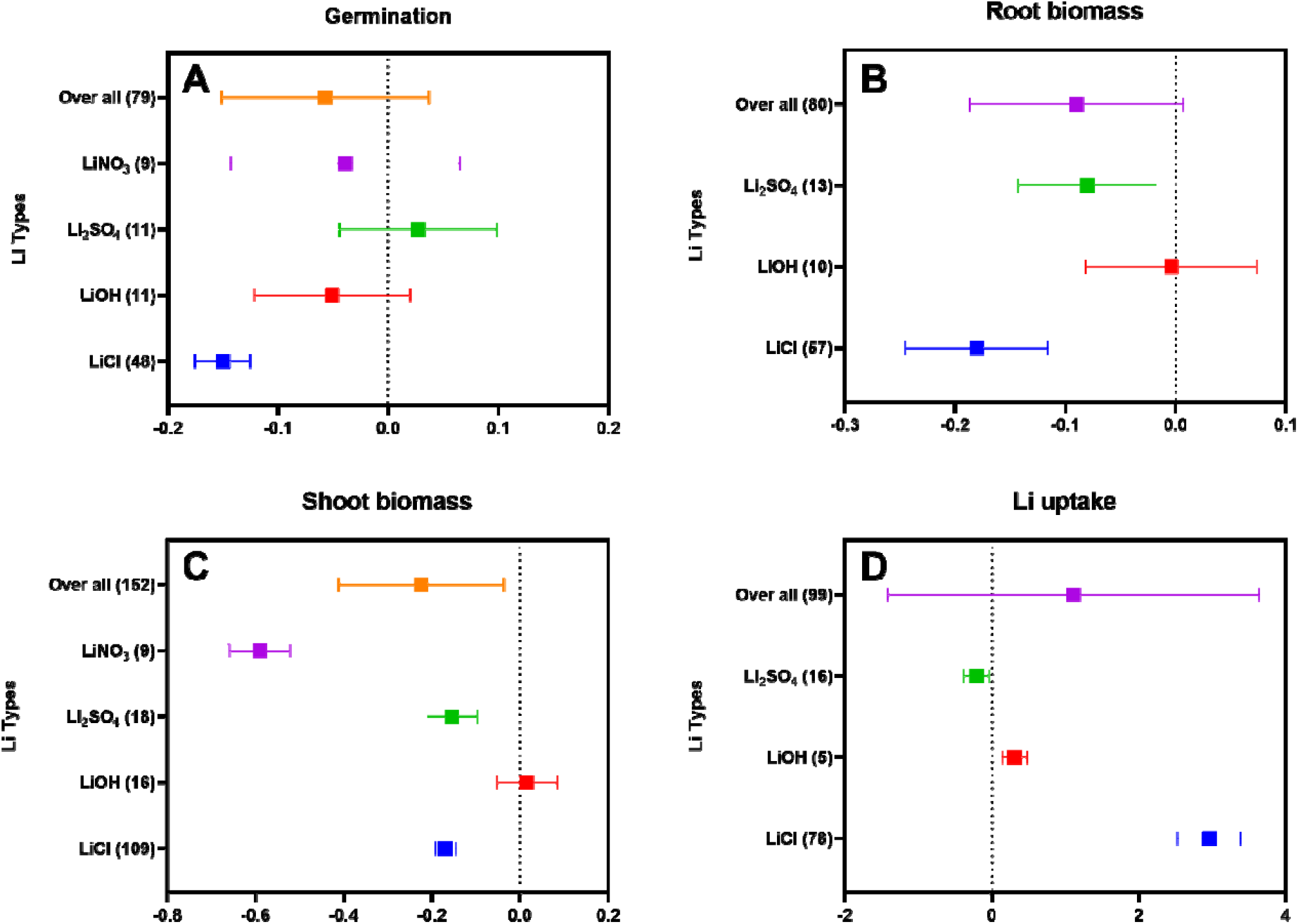
The mean effect of Li sources on germination (A), root biomass (B), shoot biomass (C) and Li uptake (D). The lift side bars presented the Li sources, and the corresponding number of endpoints is shown within the parentheses.

### 3.2 Impacts of different Li sources on physiological parameters of plants

In terms of Li sources, plant response to LiCl exposure was studied more frequently than LiOH, Li_2_SO_4_ and LiNO_3_. As measured endpoints, 272 (71%) focused on LiCl compared to 42 (10%), 47 (11%) and 18 (4%) endpoints in LiOH, Li_2_SO_4_ and LiNO_3_ respectively (Figure 4). The meta-analysis results reveal that germination, root biomass, shoot biomass and Li uptake were significantly impacted by Li sources; specifically, LiCl significantly reduced germination, root and shoot biomass by approximately 14%, 16% and 16% respectively. Exposure to LiOH decreased the germination by 5%, while no effect was observed on root and shoot biomass. Application of Li_2_SO_4_ has no impact on germination, while declined the root biomass by 8% and shoot biomass was 14%. The germination (4%) and shoot biomass (45%) was negatively affected by LiNO_3_, while root biomass was not evident yet. Average across the different Li sources, the plants germination, root and shoot biomass were significantly decreased by 6%, 9% and 20% respectively. The Li uptake by plant was different under different Li sources i.e., maximum Li uptake was observed under LiCl (2331%), followed by LiOH (42%) and Li_2_SO_4_ (22%).

Soybean exposed to LiOH at 20, 40 and 60 mg dm^-3^ have reduced germination by 10, 4 and 8% respectively, while exposure of LiSO_4_ have increased germination by 13, 15 and 25% respectively.^26^ The exposure of LiCl significantly declined the root biomass (31%) of *Zea mays* at 50 mg dm^-1^.^24^ Similarly the application of LiSO_4_ declined the *Lactuca sativa L*. root biomass by 1 and 10% at 30 and 40 mg dm^-1^ respectively. In contrast, LiOH increased the *Lactuca sativa L*. root biomass by 41 and 4% at 30 and 40 mg dm^-1^ respectively. ^13^ Similarly LiCl exposure reduced the shoot biomass of sunflower and *Zea mays* by 27 and 32% respectively at 50 mg dm^-1^.^24^ In general, the meta-analysis results indicated that the Li sources effect on plant germination, root biomass, shoot biomass and Li Uptake were different with different Li types. In case of plant germination and root biomass LiCl have more toxic effect on plants. Conversely, shoot biomass was decreased with LiNO_3_. In addition, uptake of LiCl was more than LiOH and Li_2_SO_4_.

### 3.3 Impacts of Li concentrations on plants health

In general, Li concentration (n=83) had a negative effect on plant germination. Current meta-analysis findings indicated that Li significantly inhibited germination 17% at <50 ppm and 7% at ≥ 50 ppm (Figure 5A). Plant root biomass (n=88) was significantly negatively affected (10%) by Li concentration. At <50 ppm concentrations (n=46), root biomass exhibited a highly negative response by 12% reduction, while at ≥ 50 ppm concentration (n=42) values were less effected by causing 9% reduction (Figure 5B). Exposure to Li showed negative effects on shoot biomass (n=178) as a function of concentration (Figure 5C). Our meta-analysis findings indicated that shoot biomass increased by 20% with exposure of Li at >50 ppm (n=86), while reduced by 31% and 18% at 50-500 ppm (n=49) and >500 ppm (n=43) respectively.

**Figure 5.**
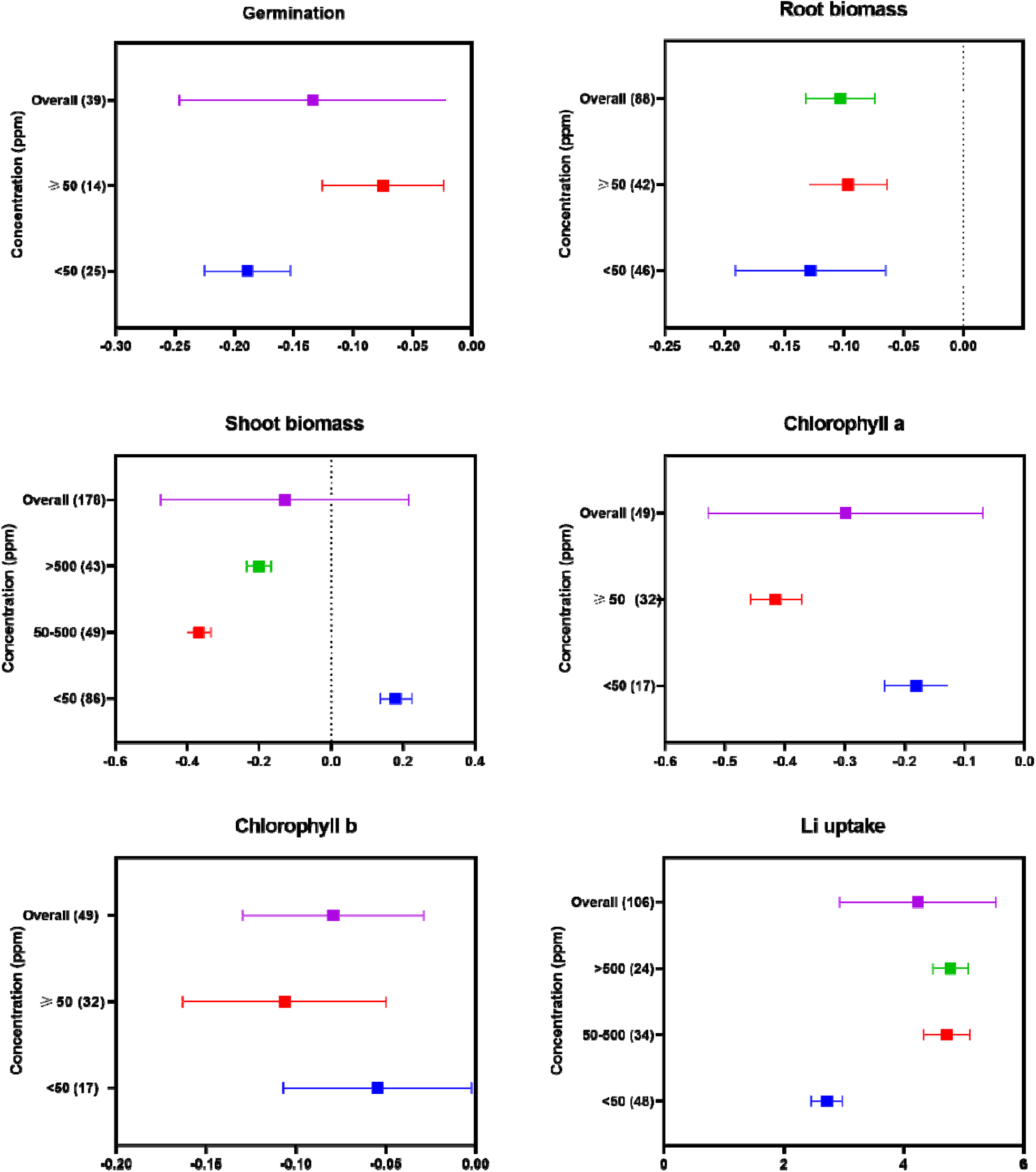
**The** mean impact of Li concentration on germination (A), root biomass (B), shoot biomass (C), chlorophyll a (D), chlorophyll b (E) and Li uptake (F). The applied concentrations are presented om lift side bars, and the corresponding number of endpoints is shown within the parentheses.

Chlorophyll a and b contents were significantly affected due to different concentrations of Li (Figure 5D and E). Chlorophyll a and b decreased by 17% and 5% at <50 ppm concentration with exposure at ≥ 50 ppm concentrations declined chlorophyll a and b by 34% and 10% respectively. The Li uptake (106) by plant was significantly high at >500 ppm (n=24) followed by medium concentration 50-500 ppm (n=34) and low concentration at <50 ppm (n=48). Overall, Li concentrations on plant germination, root biomass, shoot biomass, chlorophyll a and b were negatively affected (13%, 10%, 12%, 26% and 8% respectively). In general, the current meta-analysis results indicated that physiological parameters were negatively impact with concentration of Li, excluding chlorophyll a and b which was increased with >500 mg L^-1^ concentration. Uptake of Li were increased at different concentration.

Previous study which reported that LiCl toxic effect on seed germination (*Brassica carinata*) increase with dose trend e.g., < 30 mM had a non-significant impact on seed germination whereas >30-210 mM significant (40-95%) germination inhabitation was observed as compared to control ^25^. Similar trend was observed in the germination of *apocynum pictum*, showed non-significant effect on low doses (25 mmol L-1) however significant reduction (25-95%) were observed with increasing concentration level of Li (50-400 mmol L^-1^)^15^. As well as same trend were observed in the germination of soybean under Li stress^26^. In the context of shoot and root biomass of butterhead lettuce (*Lactuca sativa* var. *capitata*) showed non-significant increment were observed at low concentration (2.5 mg dm^-3^) of Li (LiCl and LiOH) whereas, at higher concentration (>50 mg dm^-3^) of Li (LiCl and LiOH) significantly reduced the root (36 and 21%) and shoot (58 and 69%) biomass as compared to controls plant ^27^. Another study reported that shoot biomass of *Z. mays* decreased (10-87%) with increasing dose trend (40-400 ppm) than that control plant, EC_50_ value of Z. mays for plant biomass were observed 118 ppm ^28^. Plant root and shoot biomass are linked with the accumulation of Li in plant body e.g., at higher concentration accumulating of Li increased with exogenous level that resulting reducing the plant shoot and root biomass ^28, 29^. A previous study documented that when the Li concentration level in leaves of the plant reaches a threshold level, that causes the necrosis on the older leaves, leading to the reduction of chlorophyll content of plants, ultimately reducing the plant shoot and root biomass ^27^.

Li has significant reduction effect on the chlorophyll a and b at higher Li concentration. Such as, chlorophyll a and b content of *Brassica carinata* significantly reduced at 60 mM concentration and higher reduction effect were observed with increasing dose levels, whereas non-significant reduction noticed at lower dose (0.03-30 mM) of LiCl ^25^. Another study reported that the content of chlorophyll a and b remarkably reduced at 20 mg kg^-1^ but at higher concentration (40-80 mg kg^-1^) showed non-significant impact as respective to non-treated plant^18, 23^. So, chlorophyll content depends on the plant species and dose level applied Li. The bioaccumulation of Li in plant tissues (shoot and root) significantly increased with exogenous Li concentration for example, Li concentrations in lettuce plant shoot and root increase with dose level (20-80 mg kg^-1^) as respective to control plant^18^. The accumulation of Li in plant tissues was several hundred greater than that applied concentration^18, 27^. Li is highly mobile from root and shoot and its translocation factor increasing with dose levels in the lettuce plant^27^. The accumulation of Li depend on plant species and plant parts ^30^ such as, highest amount of Li accumulated in bottoms leaves followed by the middle leaves, shoot, top leaves, roots and flowers of the distribution in different parts of plant (*Beta vulgaris* L.)^29^. Subsequently, Li accumulation in the plant tissues depend on the exogenous level of Li.

## Research gaps and perspective

Our current meta-analysis results show that Li affected a broad range of endpoints from over 50 studies and suggest general toxicity based on physiological indicators. Importantly, the literature is both limited and highly variable in terms of Li sources making the existing data somewhat inadequate to fully assess the risk of Li exposure. Our meta-analysis reveals that prior studies were conducted in soil (10%), Foliar (80%) and hydroponic (5%), these reasons add up and we are unable to draw any conclusive results regarding real environmental scenarios. Based on our meta-analysis of what has been published, the following topics are recommended for future research work:

1. Lithium impacts on soil physio chemical properties and the soil microbial communities, as well as the symbiotic relationship between soil borne microbes and plants needs serious attention. Because plant development significantly depends on soil biota and their diversity. These assessments should include chemical interactions and the subsequent impacts on soil biota.
2. A more thorough understanding of the dose-response relationship is needed to understand short- and long-term influence, as well as the fate, adsorption and transport of Li in environmental compartments. In addition, exposure medias must be systematically evaluated and categorized in terms of risk via soil, hydroponic and foliar routes.
3. A more thorough understanding of the mechanisms of Li phytotoxicity is needed through the use molecular levels in-depth studies. Importantly, transcriptomic, proteomic and metabolomics endpoints should be measured over time so as to understand the dynamics of plant response and resilience.
4. In the context of Li containing waste management, it is crucial to develop sustainable remediation technologies as Li waste will exponentially increase in coming years. The phytoremediation of Li by using hyperaccumulator selected through bioprospecting soil ecosystems can be a sustainable and long lasting option, particularly for scenarios where the Li is high and associated with co-contaminants.

It is also important to note that much of the existing literature includes studies that have doses orders of magnitude beyond that expected in the environment. We need to focus on generating hard data that may increase our understanding of exposure and risk under conditions of environmental relevance. Solid waste management, remediation, green chemistry, and circular economy are all independent fields but their integration with regulatory support will be helpful to reduce the emerging burden of Li in the natural environment.

