## Supplementary Data for "Responses of Agricultural plants to Lithium pollution: Trends, Meta-Analysis, and Perspectives"

Noman Shakoor^1†^, Muhammad Adeel^2^*^†^, Imran Azeem^1†^, [Muhammad Arslan Ahmad](https://www.sciencedirect.com/science/article/pii/S0166526X21000131?via%3Dihub#!)^3^, [Muhammad Zain^4^](https://www.sciencedirect.com/science/article/pii/S0301479720303005?via%3Dihub#!), Aown Abbas^5^, [Pingfan Zhou^1^](https://www.sciencedirect.com/science/article/pii/S0304389421005379?via%3Dihub#!), [Yuanbo Li](https://www.researchgate.net/profile/Yuanbo-Li-5?_sg%5B0%5D=mc55MYmRA3g7fNXMI6Qqgz3QsqamnOmXG-M6ZGFm-rHHuK7M7qrkGTATSp_WA-eudS8F-eA.E7lJvrAfhczbupMJLOc35xLlhFjU_zpLu-NpC7K3sqBhZvLJwY6VkcWheWW81DtBSyx-H__ChgJDOv0XxBHkHQ&_sg%5B1%5D=uZZB98pxc7XYVncSgk2qOleWzvN5zU5jzRn-j-rtSvhLI2P5RFYS54dOIcw0hETSG2438IY.FFdAQCejmiGsAPC4vzfB6YUQCDeLovJ9f-bmOM_3WvBmXpwkSBp270--c_gXWg2mPyRAznSHd2xPDHBmT-G_Cw)^1^, Xu Ming^2^ and [Yukui Rui](https://www.sciencedirect.com/science/article/pii/S0304389421005379?via%3Dihub" \l "!)^[1](https://www.sciencedirect.com/science/article/pii/S0304389421005379?via%3Dihub" \l "!)^*

^1^Beijing Key Laboratory of Farmland Soil Pollution Prevention and Remediation and College of Resources and Environmental Sciences, China Agricultural University, Beijing 100193, PR China

^2^BNU-HKUST Laboratory of Green Innovation, Advanced Institute of Natural Sciences, Beijing Normal University at Zhuhai, 18 Jinfeng Road, Tangjiawan, Zhuhai, Guangdong

^3^College of Physics and Optoelectronic Engineering, Shenzhen University, Shenzhen 518060, China

^4^Key Laboratory of Crop Water Use and Regulation, Ministry of Agriculture and Rural Affairs, Farmland Irrigation Research Institute, Chinese Academy of Agricultural Sciences (CAAS), Xinxiang, Henan, 453003, PR China

^5^Department of soil and climate change, The university of Haripur, 22780, Pakistan

*Corresponding authors:

Yukui Rui:

Muhammad Adeel:

†These authors contributed equally to this work.

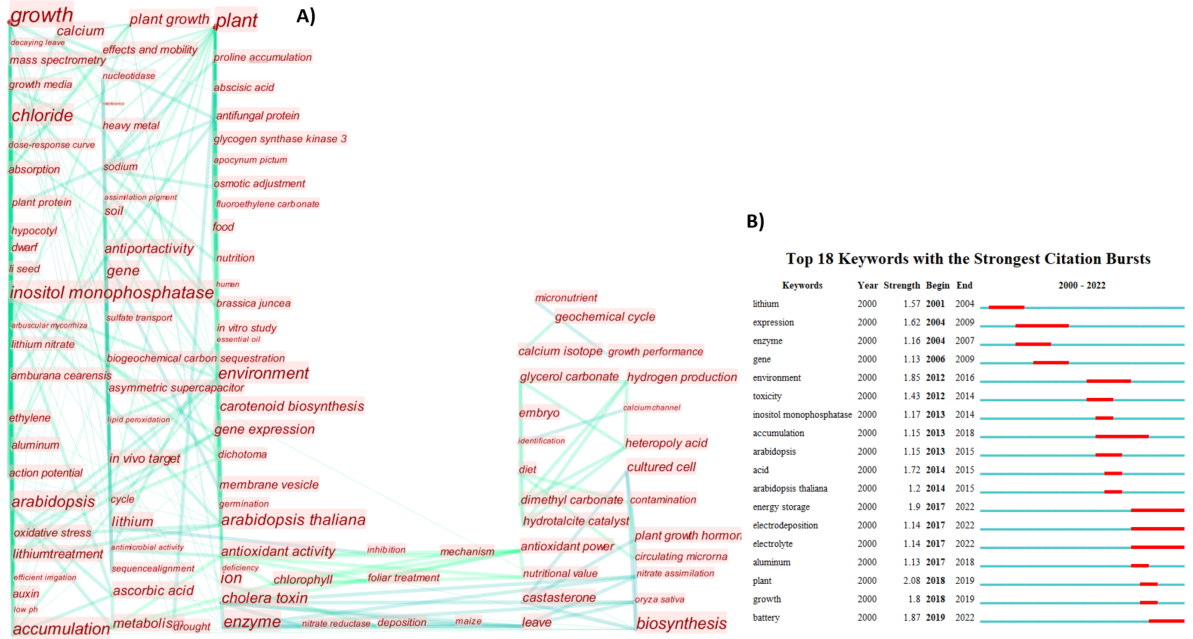

Figure S1. The chart of high frequency keywords with time and keywords clustering from 2000 to March 2022. The size of node indicates the frequency of keyword occurrence in the studied documents. (B) The top 18 keywords with the highest citation activity.
